# Low-frequency Neural Activity Reflects Rule-based Chunking during Speech Listening

**DOI:** 10.1101/742585

**Authors:** Nai Ding, Peiqing Jin

## Abstract

Cortical activity tracks the rhythms of phrases and sentences during speech comprehension, which has been taken as strong evidence that the brain groups words into multi-word chunks. It has prominently been argued, in contrast, that the tracking phenomenon could be explained as the neural tracking of word properties. Here we distinguish these two hypotheses based on novel tasks in which we dissociate word properties from the chunk structure of a sequence. Two tasks separately require listeners to group semantically similar or semantically dissimilar words into chunks. We demonstrate that neural activity actively tracks task-related chunks rather than passively reflecting word properties. Furthermore, without an explicit ‘chunk processing task,’ neural activity barely tracks chunks defined by semantic similarity - but continues to robustly track syntactically well-formed meaningful sentences. These results suggest that cortical activity tracks multi-word chunks constructed by either long-term syntactic rules or temporary task-related rules. The properties of individual words are likely to contribute only in a minor way, contrary to recent claims.

## Introduction

How the brain processes language is a central question in cognitive science, neuroscience, and linguistics. In general, speech utterances such as sentences are not memorized sequences, but instead are proposed to reflect compositional processes which allow us to understand and produce countless new sentences never heard before (such as the one you are currently reading). Therefore, to derive the meaning of a sentence, the brain has to integrate information across words, the meaning of which are stored in long-term memory. How the brain integrates information across words, however, remains debated (Hagoort and Indefrey, 2014; Goucha et al., 2017; Romeo et al., 2018), and a variety of hypotheses have been proposed (Townsend and Bever, 2001; Ferreira et al., 2002; Everaert et al., 2015). At one end of the spectrum, it has been hypothesized that the brain applies a set of syntactic rules to recursively combine words into larger chunks, forming a hierarchically organized syntactic structure, and then derives meaning of a sentence based on its syntactic structure (Chomsky, 1957; Frazier and Fodor, 1978; Friederici, 2002). At the other end of the spectrum, it has been hypothesized that the brain does not construct multi-word chunks at all but instead directly integrates information across words by statistical and semantic analysis (Elman, 1990; Frank et al., 2012; Christiansen and Chater, 2016). It is challenging to adjudicate among these hypotheses, as multi-word chunks are ‘intermediate’ mental representations that can are hard to measure directly.

Recent neurophysiological results have been shown that when presented with speech, the brain concurrently tracks multiple levels of linguistic units, such as sentences, phrases, words, and syllables (Fig. 1) (Ding et al., 2016; Makov et al., 2017; Brodbeck et al., 2018; Broderick et al., 2018; Ding et al., 2018; Keitel et al., 2018). Critically, neural tracking of phrases and sentences remains when the phrasal and sentential boundaries are not cued by prosodic features or by the transitional probability between words. This phenomenon has been taken as strong evidence for the hypothesis that the brain applies grammatical rules to group words into chunks (Ding et al., 2016; Martin and Doumas, 2017). This hypothesis is referred to as the rule-based chunking hypothesis in what follows. Challenging this position, it has been argued that neural tracking of phrases and sentences can potentially be explained by neural tracking of properties of individual words alone. In the following, we illustrate these ideas using sentence-level tracking as an example; the same principle also applies to phrase-level tracking.

**Figure 1.**
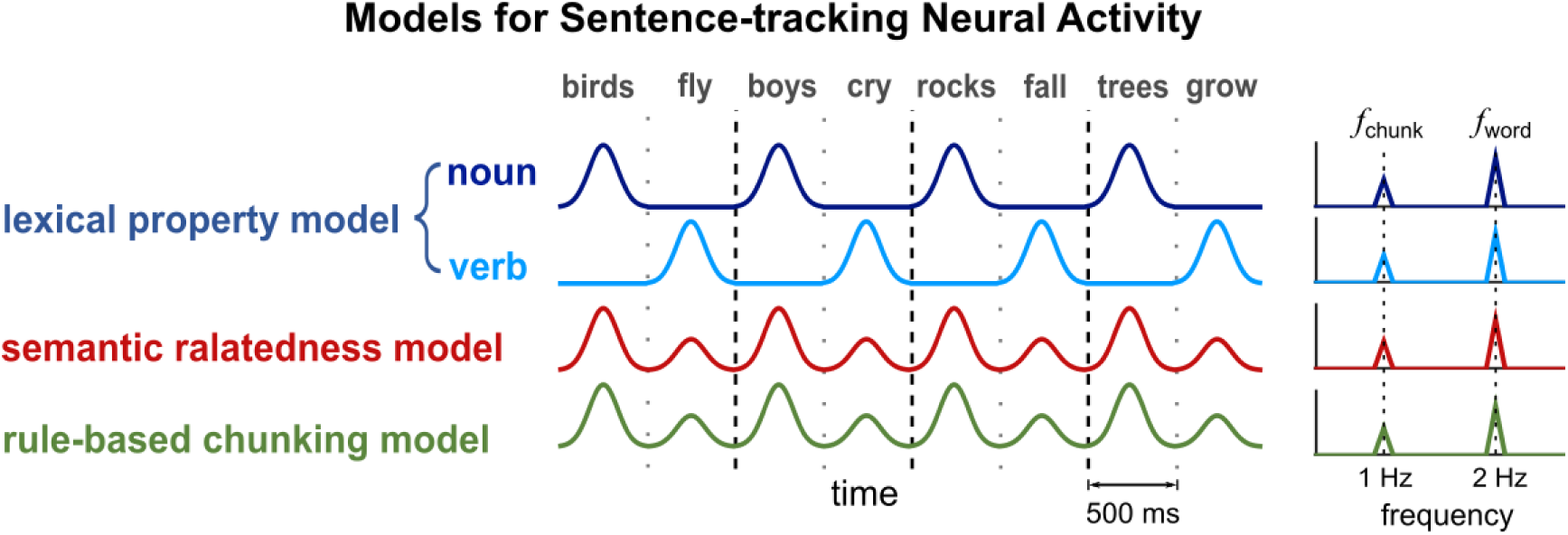
Three models for sentence-tracking neural activity. The stimuli in this illustration are all 2-word sentences (noun + verb). Words are isochronously presented at 2 Hz. Sentence boundaries are marked by black dashed lines while word boundaries within a sentence are marked by gray dotted lines. The lexical property model assumes neurons that are selectively tuned to nouns and verbs. The semantic relatedness model assumes that the neural response to a word is attenuated if the word is preceded by a semantically related word. Additionally, it is assumed that words within a sentence are more closely related than words across a sentence boundary. The rule-based chunking model assumes a consistent change of neural activity within a mentally constructed chunk. Here, to facilitate the comparison with the semantic relatedness model, it is further assumed that the neural response is stronger at the chunk onset. All the 3 models can generate sentence-tracking neural activity.

First, it is known that some neural populations are selectively tuned to words from specific syntactic (Caramazza and Hillis, 1991; Daniele et al., 1994) or semantic categories (Warrington and Shallice, 1984; Bi et al., 2016). For the experimental paradigm shown in Fig. 1, all sentential stimuli have the same syntactic structure. Therefore, neural activity tracking lexical properties, such as part of speech information or lexical semantic information that distinguishes objects and actions, may appear to track sentences (dark and light blue curves in Fig. 1). Consistent with this lexical property model, it has been shown that apparent neural tracking of sentences could be observed if the neural response independently represents each word using multidimensional features learned by statistical analysis of large corpora (Frank and Yang, 2018). In other words, neural populations that are tuned to lexical features of individual words may show activity that apparently tracks sentence structures.

Second, it is well established that the neural response to a word depends on the context and the response amplitude is smaller if the word is semantically related to previous words (Lau et al., 2008; Kutas and Federmeier, 2011). In general, words within the same sentence are more related than words from neighboring sentences. Therefore, for a context-dependent neural response, its amplitude is expected to be stronger at the beginning of a sentence, leading to apparent neural tracking of sentences (red curve, Fig. 1). This model considers semantic relatedness between words, but it does not consider the sentence structure: Semantic relatedness is evaluated the same way within and across sentence boundaries. Apparent sentence tracking behavior is generated since words within a sentence are more closely related.

The lexical property model and semantic relatedness model do not assume linguistic chunks and therefore provide different explanations for sentence/phrase-tracking responses than the rule-based chunking model (green curve, Fig. 1). The rule-based chunking model, however, has additional flexibility, allowing the same sequence of words to be grouped differently based on different sets of rules. This flexibility is most clearly demonstrated when processing structurally ambiguous sequences. For example, “sent her kids story books” can be chunked as “sent [her kids] story books” or “sent her [kids story books]” (Shultz and Pilon, 1973). For such structurally ambiguous sentences, the rule-based chunking model, but not the lexical property model or the semantic relatedness model, would predict different phrase-tracking responses when the sentence is chunked differently.

Here, we distinguish the rule-based chunking model from the word-based models by designing a structurally ambiguous word sequence and two different rules to bias the chunking of this sequence. The word sequence is a sequence of nouns that describe either living (L) things or nonliving (N) things (Fig. 2A). The sequence has no syntactically defined chunks. Nevertheless, the participants are explicitly instructed about how to chunk the sequence, and the chunks are differently constructed in 2 separate conditions. While participants listen to the word sequences, we recorded cortical responses using MEG. Both word-level models, i.e., the lexical property model and the semantic relatedness model, predict the same neural response when participants listen to the same word sequence regardless of how the sequence is chunked. The rule-based chunking model, however, predicts chunk-dependent neural responses. Additionally, we also tested how the brain encodes the word sequence and sentence sequences without explicit chunking tasks.

**Figure 2.**
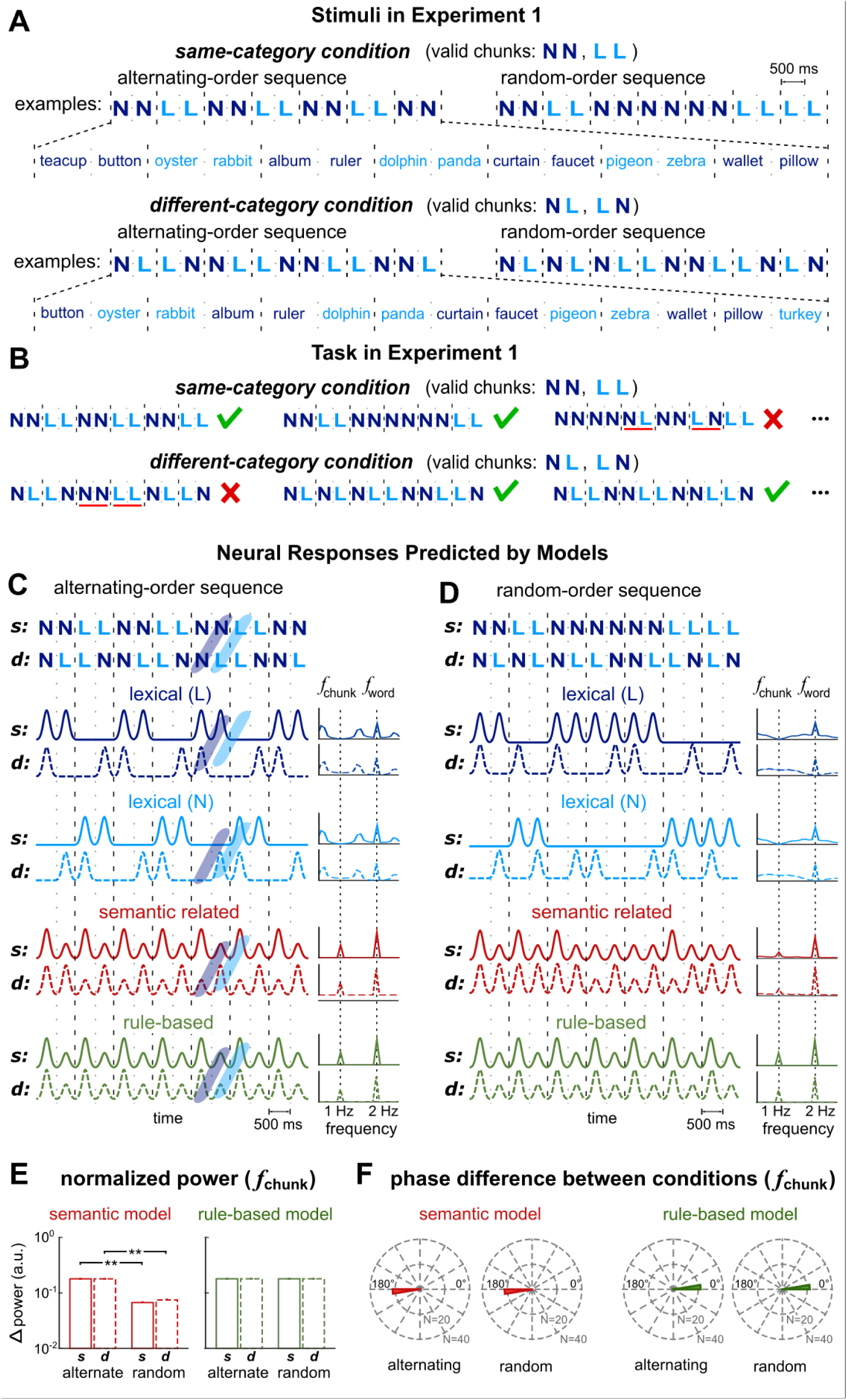
Stimuli in Experiment 1 and model simulations. (A) Stimuli consist of isochronously presented nouns, describing either living (L) or nonliving (N) things. Two words construct a chunk and the chunks further construct sequences. Different chunks are used in the same-category condition (upper panel) and different-category condition (lower panel). Sequences in each condition further divides into alternating-order (left panel) and random-order (right panel) sequences. In the illustration, chunk boundaries are marked by black dashed lines while word boundaries within a chunk are marked by gray dotted lines. (B) The task is to decompose each sequence into 2-word chunks and detect invalid chunks. Three trials and the correct responses are shown for each condition (tick and cross for normal and outlier trials respectively). Red underlines highlight the invalid chunks. (C) Predicted neural responses to the alternating-order sequences. The lexical model separately considers neurons selectively tuned to living and nonliving things. The semantic relatedness model and the rule-based chunking model, but not the lexical property model, generate a chunk-rate response. The tilted blue regions illustrate that the alternating-order sequences only differ by a time lag between the same-category and different-category conditions. Neural responses predicted by the lexical property and semantic relatedness models also differ by a time lag between conditions. (D) Predicted responses to the random-order sequences. (E) Predicted chunk-rate response power. The semantic relatedness model, but not the rule-based chunking model, predicts a difference in response power between alternating- and random-order sequences. (F) The semantic relatedness model predicts a 180° phase difference between same- and different-category conditions, while the rule-based chunking model predicts a 0° phase difference. * P <0.05, ** P < 0.01

## Results

### Word Sequences and Model Predictions

In Experiment 1, participants were instructed to parse a sequence of words into chunks and each chunk consisted of 2 words. The words were drawn from 2 categories, i.e., living and nonliving things. The experiment contrasted 2 conditions in which the chunks were constructed based on different rules. In one condition, referred to as the same-category condition, the 2 words in a chunk belonged to the same semantic category (Fig. 2A, upper panel). In the other condition, referred to as the different-category condition, the 2 words in a chunk were drawn from different categories (Fig. 2A, lower panel). Based on these rules, there were 2 valid chunks in the same-category condition, i.e., LL and NN, and also 2 valid chunks in the different-category condition, i.e. NL and LN.

The same- and different-category conditions were presented in separate blocks. In each condition, participants had to parse the sequences into chunks, i.e., pairs of words, and detect invalid chunks that were occasionally presented. They had to press different keys after trials containing or not containing invalid chunks (Fig. 2B). The sequences in each condition (*N* = 60) were further divided into 2 types of sequences, i.e., the alternating-order sequence (*N* = 30) and the random-order sequence (*N* = 30). In the alternating-order sequence, the 2 valid chunks in each condition were interleaved while in the random-order sequence the 2 valid chunks were presented in random order. The alternating-order sequences and the random-order sequences were intermixed and presented randomly in each block. The neural responses to these 2 types of sequences were separately analyzed. The alternating-order sequences allowed an intuitive comparison between the same- and different-category conditions, which would be detailed in the following. The random-order sequences were designed as fillers to increase variability, but model simulations in the following show that they can also distinguish the 3 hypotheses in Fig. 1.

Simulations of the neural responses to the alternating- and random-order sequences by the 3 models are shown in Fig. 2 for both the same- and different-category conditions. The lexical property model considers 2 neural populations that selectively respond to living and nonliving word meanings, respectively. The simulated neural response analyzed in the frequency domain demonstrates that none of the population shows a chunk-rate response peak in the spectrum (dark and light blue curves in Fig. 2CD). In contrast to the lexical property model, the semantic relatedness model and the rule-based chunking model predict a chunk-rate response peak (red and green curves respectively in Fig. 2CD). The semantic relatedness model further predicts that the chunk-rate response is stronger for the alternating-order sequence than the random-order sequence (Fig. 2E). The rule-based chunking model, however, predicts similar response amplitude for the 2 kinds of sequences (Fig. 2E).

A more fundamental difference between the semantic-relatedness model and rule-based chunking model lies in their predictions about the chunk-rate response phase. The semantic-relatedness model predicts a 180° phase difference between same- and different-category conditions, while the rule-based chunking model predicts a 0° phase difference between conditions. For the alternating-order sequence, these predictions are straightforward: These sequences are offset by 1 word between the same- and different-category conditions. Consequently, neural activity tracking semantic relatedness between words is offset by the duration of a word between conditions. For neural activity oscillating at the chunk rate, this time lag leads to a 180° phase difference (Fig. 2F). For the random-order sequence, although less straightforward, model simulation shows that the chunk-rate response has a 180° phase difference between conditions (Fig. 2F). For the rule-based model, however, the response is aligned with the chunk boundaries, which are not affected by the conditions and sequence types. Therefore, the rule-based model predicts the same response phase, i.e. a 0° phase difference, for the same- and different-category conditions (Fig. 2F).

In summary, the 3 models considered in this study lead to different predictions about the neural response properties (Fig. 2). Details about the model simulations are given in Supplementary Fig. 1. The lexical property model and the semantic relatedness model are simulated based on neurons that have ideal tuning to living/nonliving things, but similar results can be obtained using realistic semantic models of words, i.e., the word2vec model learned by statistical analysis of natural language (Supplementary Fig. 2). The word2vec model, a connectionism model that does not explicitly model phrasal structures, can successfully capture semantic relationship between words (Bengio et al., 2003). In the following, we turn to the actual neural responses obtained using MEG and evaluate their consistency with the simulations made for the 3 different models.

### Task-dependent Neural Tracking of Alternating-order Sequences

First, we analyzed the MEG responses to the alternating-sequences. The MEG responses were separately averaged for the same- and different-category conditions and the mean response was transformed to the frequency domain. The response spectrum averaged over all MEG gradiometers showed clear peaks at the chunk and word rates (Fig. 3A, left 2 plots). The chunk-rate response power was significantly stronger than the power in neighboring frequency bins in both conditions (P = 0.0001 for both the same- and different-category conditions; bootstrap, Methods, FDR corrected). The chunk-rate spectral peak was consistent with the semantic relatedness model and the rule-based chunking model, but not with the lexical property model (Fig. 2C). More importantly, the phase difference between conditions is closer to 0° than 180° in all MEG sensors (Fig. 4AB), consistent with the rule-based chunking model (Fig. 2F). Specifically, the response phase difference was significantly different from 180° in 99% of the sensors (303 out of 306, P < 0.01, bootstrap, Methods, not corrected for multiple comparisons). For the response phase difference averaged over all MEG sensors, the 99% confidence interval over participants ranged from −50° to 15°, which is centered around 0° and does not include 180°.

**Figure 3.**
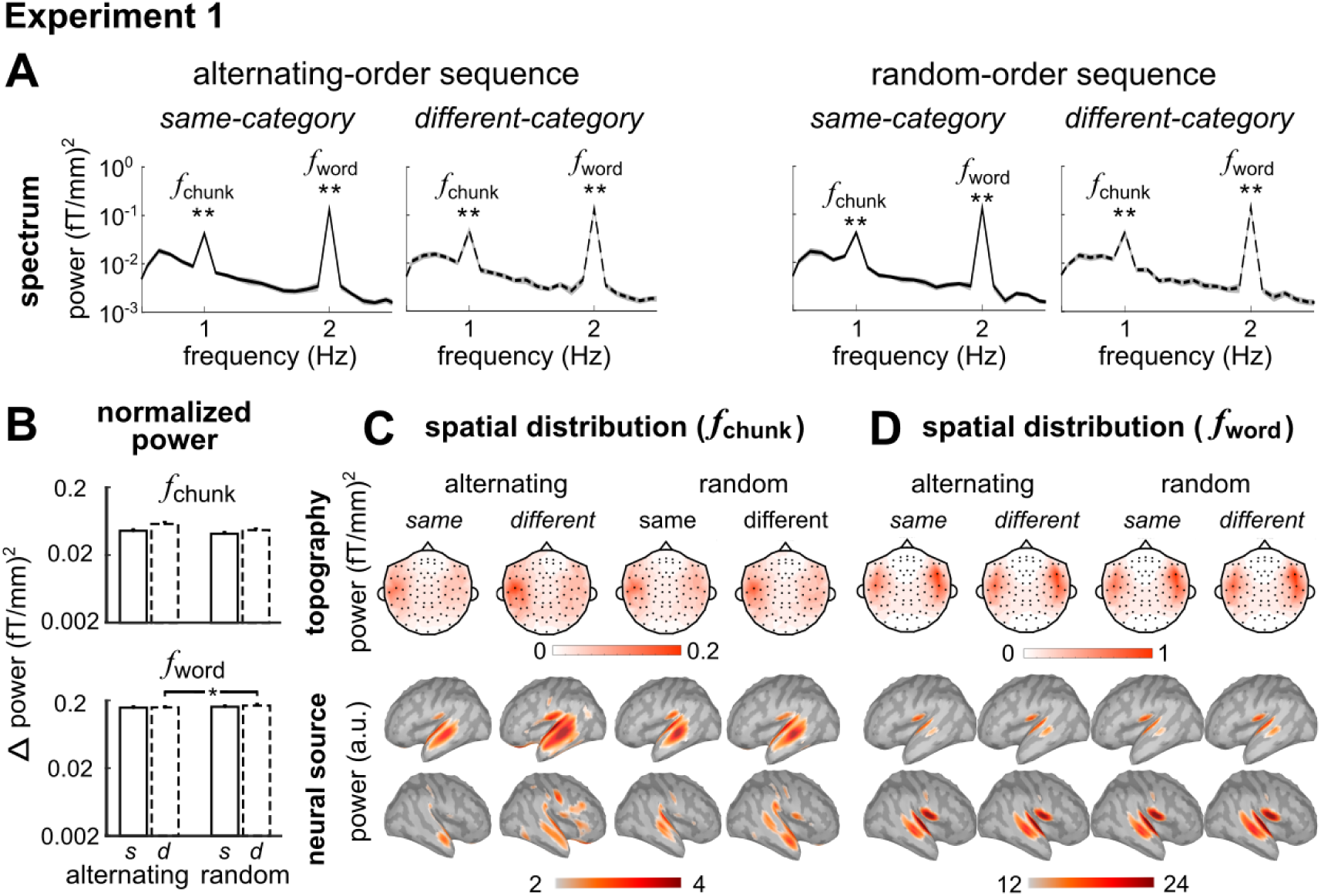
Response power results of Experiment 1. (A) The response power spectrum averaged over participants and MEG gradiometers. A chunk-rate response peak and a word-rate response peak are observed. The shaded area covered 1 SEM over participants on each side. (B) Normalized chunk- and word-rate response power. The chunk- and word-rate response power does not significantly differ between conditions. (C, D) Response topography (gradiometers) and source localization results, averaged over participants. Only statistically significant sensors (shown by black dots) and vertices are shown (P < 0.05, FDR corrected) are shown in the topography and localization results. Chunk-rate and word-rate responses are mainly generated from bilateral temporal areas. The neural source localization results are shown by the dSPM values. * P <0.05, ** P < 0.01

In the response topography, the chunk- and word-rate responses both show bilateral activation over the temporal lobes (Fig. 3CD). The chunk-rate response was stronger in the left hemisphere: The response averaged over left hemisphere sensors was significantly stronger than the response averaged over the right hemisphere sensors (P = 0.014 and 0.017 for the same- and different-category conditions respectively; bootstrap, Methods). The word-rate response was significantly stronger in the right hemisphere for the same-category condition (P = 0.019; bootstrap, Methods) but did not differ across the different-category condition (P = 0.157; bootstrap, Methods). Neural source localization showed that both the chunk- and the word-rate responses were mainly generated from bilateral temporal lobes (Fig. 3CD).

### Task-dependent Neural Tracking of Random-order Sequences

The MEG responses to random-order sequences was analyzed the same way as the responses to alternating-order sequences. In the response spectrum, clear peaks were observed both for the chunk and word rates (Fig. 3A, right 2 plots). The chunk-rate response power was significantly stronger than the power in neighboring frequency bins in both conditions (P = 0.0001 for both the same- and different-category conditions; bootstrap, Methods, FDR corrected). The chunk-rate response power was not significantly different between random- and alternating-order sequences (P = 0.879 and 0.320 for same- and different-category conditions respectively; bootstrap, Methods, FDR corrected; Fig. 3B). The word-rate response power was significantly stronger for random-order sequence than alternating-order sequence in different-category condition (P = 0.022; bootstrap, Methods) but not for the same-category condition. Furthermore, for the chunk-rate response, the phase difference between conditions was closer to 0° than 180° in all MEG sensors (Fig. 4DE). The phase difference was significantly different from 180° in 99% of the sensors (303 out of 306 sensors, P < 0.01, bootstrap, Methods, not corrected for multiple comparisons). For the phase difference averaged over MEG sensors, the 99% confidence interval over participants ranged from −45° to 71°, which was centered around 0° and did not include 180°. Altogether, both the power and phase results are consistent with the prediction of the rule-based chunking model. The response waveform is shown for a representative MEG sensor, which oscillates in phase in the 2 conditions (Fig. 4CF). The representative sensor has the highest normalized power at the chunk rate, when averaged over the 2 conditions.

In the response topography, the chunk- and word-rate responses both showed bilateral activation over the temporal lobes (Fig. 3CD). The chunk-rate response was significantly stronger in the left hemisphere for the different-category condition (P = 0.0002; bootstrap, Methods) but not for the same-category condition (P = 0.353; bootstrap, Methods). The word-rate response showed no significant lateralization (P = 0.103 and P = 0.089 for same- and different-category conditions respectively; bootstrap, Methods). Neural source localization results showed that both the chunk- and the word-rate responses were mainly generated from bilateral temporal lobes (Fig. 3CD).

### Task Modulation of Neural Tracking of Semantic Chunks

In Experiment 1, we designed 2 sets of chunks and explicitly asked participants to parse sequences into 2-word chunks. In Experiment 2, we further investigate whether semantic similarity can automatically drive chunk-tracking neural responses without an explicit chunking task. For this purpose, we presented sequences that had the same structure as the alternating-order sequences in the same-category condition of Experiment 1. Participants, however, perform 3 different tasks in separate blocks. The first task consisted of a chunk-level task, similar to the task in Experiment 1, i.e. detecting invalid chunks. The second task consisted of a word-level task, detecting if a concrete noun, i.e., living or nonliving things, was replaced by an abstract noun. The third task was an auditory task, detecting a change in the speaker’s voice.

Fig. 5A, B, and C show the MEG response spectrum for the chunk-level, word-level, and auditory tasks, respectively. When participants perform the chunk-level task, a strong chunk-rate response was observed in the spectrum, replicating results of Experiment 1. When performing the auditory and word-level tasks, however, the chunk-rate responses were weaker. The chunk-rate response power was significantly stronger than the power in neighboring frequency bins during all 3 tasks (P = 0.0001, 0.001, and 0.0001 for chunk-level, word-level and auditory tasks respectively; bootstrap, Methods, FDR corrected). While the chunk-rate response was statistically significant for all 3 tasks, the response was significantly stronger in the chunk-rate task than the word-level task (7.6 dB difference; P = 0.0006; bootstrap, Methods) and the auditory task (8.2 dB difference; P = 0.0004; bootstrap, Methods). These results indicate that semantic similarity between words can drive a chunk-tracking response without an explicit chunking task but the chunk-tracking response is greatly enhanced during an explicit chunking task.

In the response topography, the chunk- and word-rate responses both showed bilateral activation over temporal lobes (Fig. 5). The chunk-rate responses were stronger in the left hemisphere during the chunk-level task (P = 0.0002; bootstrap, Methods) but not during the word-level and auditory tasks (P = 0.209 and 0.158 respectively; bootstrap, Methods). The word-rate response showed left lateralization during the auditory task (P = 0.042; bootstrap, Methods) but not during the chunk-level and word-level tasks (P = 0.058 and 0.065 respectively; bootstrap, Methods). Source localization showed that both the chunk- and the word-rate responses were mainly originated in bilateral temporal lobes (Fig. 5).

### Task-irrelevant Neural Tracking of Sentences and Word Sequences

The data above show that neural activity can spontaneously track word chunks defined by semantic similarity. However, without an explicit chunking task, the chunk-rate response is rather weak. Here, a control condition is used to further validate whether the weak chunk-rate response is driven by semantic similarity, rather than, e.g., the tendency to segment any word sequence into 2-word chunks. In the control condition, living and nonliving things were presented in a random order, not consistently forming pairs belonging to the same semantic category.

Additionally, chunking of natural languages relies on both semantic and syntactic cues. Therefore, in the final condition, we investigate whether word chunks defined by syntactic rules can drive chunk-rate responses even in the absence of an explicit chunking task. In the sentence condition, each sentence started with a 2-syllable noun, followed by a 2-syllable verb. In both the control condition and the sentence condition, participants performed a word-level task, detecting occasionally occurring abstract nouns.

The responses to sentences and random words are shown in Fig. 5DE. During the word-level task, a strong chunk-rate response was observed for the sentence condition but not for the control condition. Only for the sentence condition, the chunk-rate response power was significantly stronger than the power in neighboring frequency bins (P =0.0002; bootstrap, Methods). The chunk-rate response was also stronger in the sentence condition than the control condition (P = 0.0004; bootstrap, Methods). Furthermore, the chunk-rate response was also stronger for sentences than chunks grouped by semantic similarity (P = 0.001, 0.0004, and 0.0004 for the chunk-level, word-level, and auditory tasks respectively; bootstrap, Methods; FDR corrected). Therefore, rules drive a chunk-rate response more strongly than semantic similarity.

As for the response topography, in the sentence condition, the chunk-rate response topography showed activation over bilateral temporal lobes (Fig. 5DE) and the activation was stronger in the left hemisphere (P = 0.0002; bootstrap, Methods). In the control condition, the chunk-rate response was not statistically significant, without clear activation in the sensor and source space. The word-rate response also showed activation over bilateral temporal lobes and the activation was stronger in the left hemisphere for both the sentence and control condition (P = 0.049 and 0.027 respectively; bootstrap, Methods). Neural source localization confirmed that the chunk-rate response in the sentence condition and the word-rate responses were mainly generated from bilateral temporal lobes (Fig. 5DE).

## Discussion

How words are grouped into multi-word chunks such as phrases and sentences during speech comprehension is a prominent question in psychology and cognitive neuroscience. Here we demonstrate that low-frequency neural activity can track multi-word chunks that are mentally constructed based on artificial chunking rules instead of word properties (Figs. 3 and 4). These results contradict models that assume word-level neural representations only and support a rule-based chunking model. The 3 models considered here, however, are all built on well-motivated psychological and neuroscientific evidence, and in the following we briefly review each model and discuss why the rule-based chunking model is the only one able to generate correct predictions about the neural responses.

**Figure 4.**
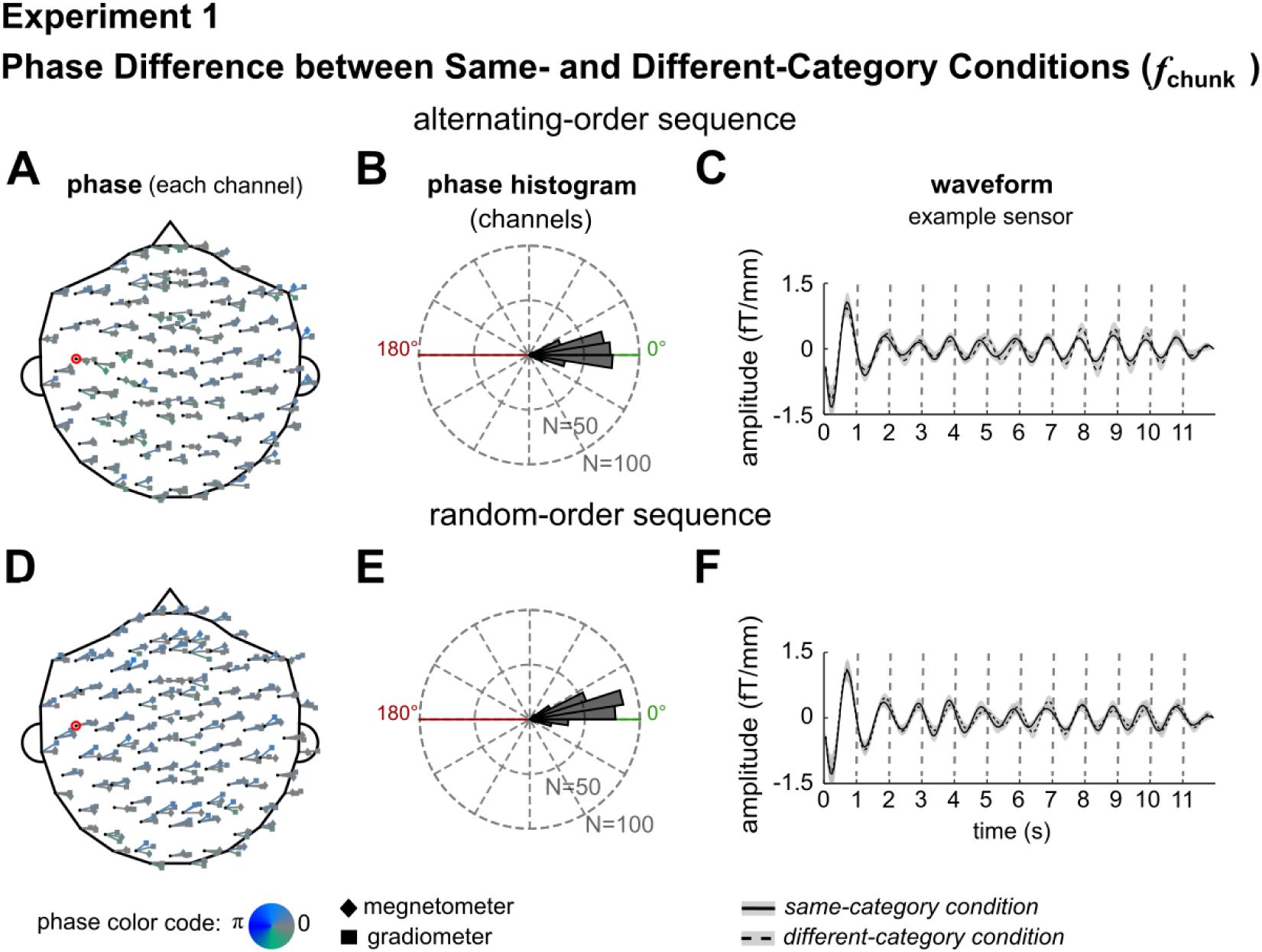
Phase difference between the same-category and the different-category conditions, for the alternating-order sequences (A-C) and random-order sequences (D-F). (A, D) Phase difference between conditions for each MEG sensor (both magnetometers and gradiometers). The results are grand averaged over participants. The phase difference angle of each sensor is indicated by a bar originating from the location of the sensor (shown by a black dot). The coherence of phase difference over participants is indicated by length of the bars. The phase difference angle is coded by both the orientation and the color of the bar. (B, E) Histogram of the phase difference for all 306 MEG sensors. The phase difference angle is closer to 0° (predicted by the rule-based chunking model) than 180° (predicted by the semantic relatedness model). (C, F) Response from a representative sensor (circled position in red in A and D). The waveform is filtered around 1 Hz, i.e., the chunk rate, which is highly consistent for the same-category chunk condition and the different-category chunk condition.

**Figure 5.**
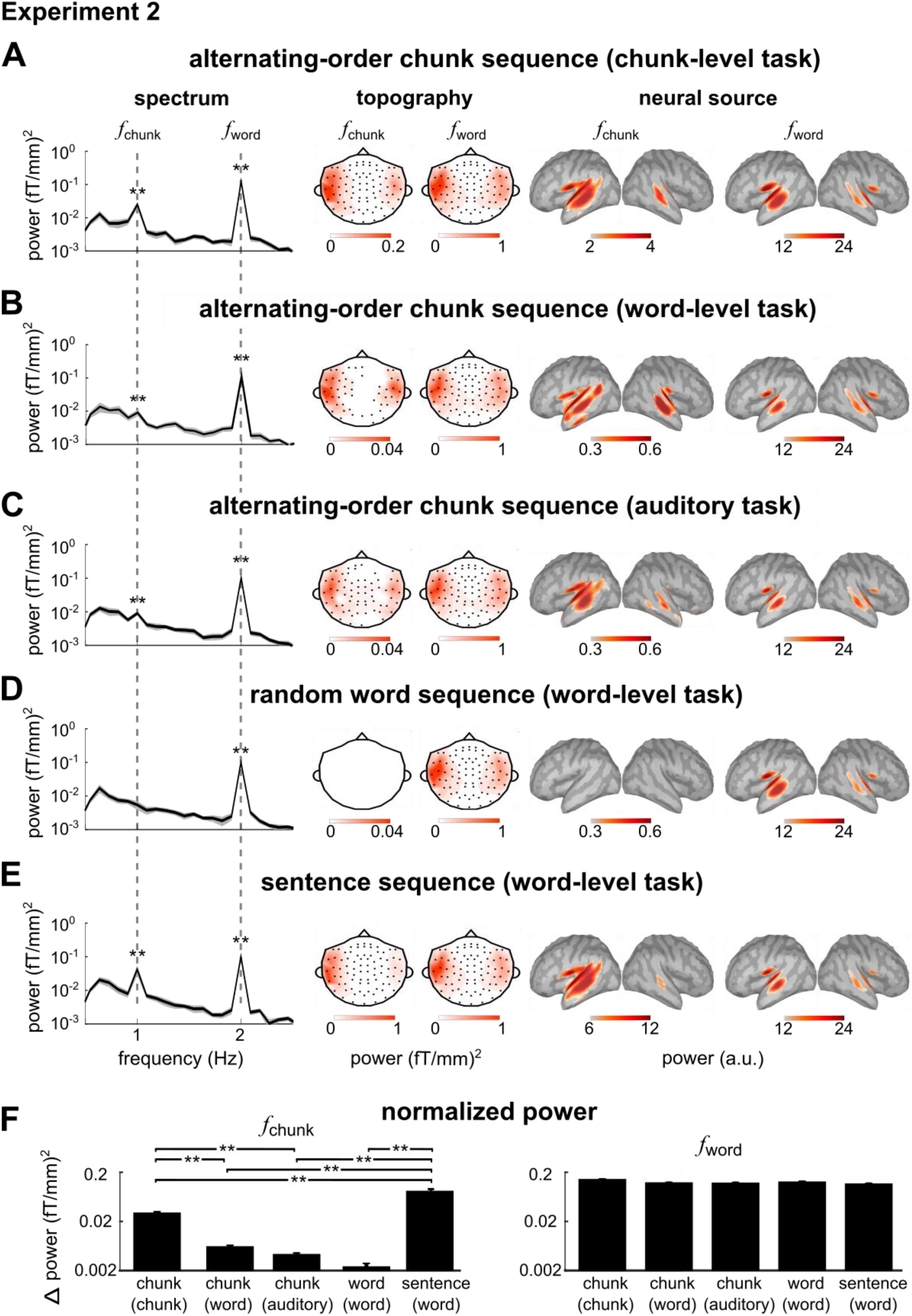
Results of Experiment 2. A-E) The response spectrum averaged over participants and MEG gradiometers are shown on the left. The response topography at the chunk- and word-rate (gradiometers) are shown in the grayscale plots in the middle. The source localization results are shown in the right (see the legend of Fig. 3 for details). For the topography and source localization results, only statistically significant sensors (shown by black dots) and vertices are shown (P < 0.05, FDR corrected) are shown. F) The normalized power at chunk and word rates. For the alternating-order chunk sequence, the chunk-rate response is stronger when participants attend to the chunks but remain significant when participants attend to words or auditory features. No significant chunk-rate response is observed for a random word sequence while a clear sentence-rate response is observed for a sentence sequence. The word-rate response is comparable across conditions. * P <0.05, ** P < 0.01

### Lexical Representation Hypothesis

The lexical representation hypothesis posits that the neural tracking of multi-word linguistic chunks is driven by neural tracking of lexical properties of individual words (Fig. 1) (Frank and Yang, 2018). This hypothesis builds on the findings that different categories of words have distinct representations in the brain. Both neurological studies and functional imaging studies have shown that words from different grammatical categories, e.g., verbs and nouns, are separately represented in the brain (Vigliocco et al., 2011; Yang et al., 2017) and can be selectively impaired (Caramazza and Hillis, 1991; Daniele et al., 1994). Similarly, objects from distinct semantic categories, e.g., living and nonliving things, are separately represented in the brain and can be selectively impaired (Warrington and Shallice, 1984; Bi et al., 2016).

Neural sensitivity to semantic categories has been mostly reported using fMRI, PET, and lesion studies, but few studies have suggested that it can be detected with the spatial resolution of MEG. For example, using a multi-channel MEG decoding technique, it has been shown that the neural responses to living and nonliving things, presented by auditory words, can be distinguished at about 70% accuracy (Chan et al., 2011). Using similar decoding approaches, the semantic categories of visually presented objects have been successfully decoded (Carlson et al., 2011; Sudre et al., 2012). Although these studies have shown that the spatial pattern of cortical activity carries semantic information using sophisticated multivariate neural decoding approaches, to our knowledge no study has shown that univariate MEG/EEG responses (e.g., single-channel responses or global field power) could clearly distinguish the semantic content of words. The neural responses to word chunks (both task-defined chunks and syntactically defined sentences), however, can be observed in single MEG/EEG sensors and in the global field power (Ding et al., 2016; Jin et al., 2018), and therefore is not easily explained by the category-dependent spatial representations in the brain.

In terms of neural sensitivity to grammatical categories of words, it is shown that the event-related potentials (ERP) evoked by verbs and nouns have statistically significant differences (Pulvermüller et al., 1999; Barber et al., 2010). This phenomenon cannot explain neural tracking of chunks of nouns but can potentially explain neural tracking of sentences. Nevertheless, the category-dependent ERP component, i.e., the difference between the ERP to different word categories, is much weaker than the ERP averaged across word categories. In other words, the EEG/MEG response evoked by an auditory word is dominated by a category-independent component. The neural response to sentences, however, is of similar amplitude to the response to words (Fig. 4), and therefore can hardly be explained by differential neural sensitivity to verbs and nouns. In sum, our study provides compelling evidence showing that words from different syntactic or semantic categories are represented by distinguishable distributive spatial patterns in the brain. For MEG/EEG responses that are summed over large-scale neural networks, however, word category information does not strongly modulate the response strength or time course and therefore barely contributes to the chunk-tracking MEG response.

### Semantic Relatedness Hypothesis and Semantic Predictions

The semantic relatedness hypothesis posits, in turn, that the neural tracking of multi-word linguistic chunks is driven by tracking of semantic relatedness between neighboring words (Fig. 1). Here, semantic relatedness refers to both semantic similarity (e.g., travel - journey) and semantic associations (e.g., travel - plan). The semantic-relatedness hypothesis builds on the priming effect in the psychological literature (Tulving and Schacter, 1990) and the neural adaptation effect in the neuroscience literature (Grill-Spector et al., 2006). It is well established that if a word is preceded by a semantically related word, its processed faster (Collins and Loftus, 1988) and its neural response, especially the ERP N400 component and its MEG counterpart, is reduced (Lau et al., 2009; Kutas and Federmeier, 2011; Broderick et al., 2018).

In this study, words from the same semantic category are more closely related than words drawn from different categories. Nevertheless, the categories used here are broad categories (e.g., animals or plants). In general, words from a broad category, e.g., animals, are only weakly related compared with words from a narrower category, e.g., birds (Vigliocco et al., 2002; Quinn and Kinoshita, 2008). A weak relationship between words predicts a weak priming effect on the neural response (Federmeier and Kutas, 1999), which may underlie why semantic relatedness between words does not explain the chunk-tracking neural activity.

The semantic relatedness model discussed here builds on the priming and neural adaptation effects. Priming and neural adaptation, however, can be caused by multiple factors and can be generally observed for any predictable stimulus (Friston, 2005; Bar, 2007; Tian and Poeppel, 2013). Previous studies have identified 2 kinds of semantic priming, i.e., automatic and strategic priming (Neely, 1977). Automatic priming can be caused by, e.g., semantic relatedness between words in long-term memory. Strategic priming, however, can actively predict upcoming words based on temporally learned association rules. Behavioral experiments have demonstrated a cross-category priming effect if the prime word from one category, e.g., tools, is known to predict target words from a different category, e.g., animals (Neely, 1977). In other words, participants can make use of association rules learned during an experiment to actively predict words that have no long-term semantic relationship with the prime word. Different from automatic priming that can occur with very short SOA between words, strategic priming occurs when the SOA between the prime and target words is relatively long, e.g., >400 ms (Hutchison, 2007).

In the current study, the SOA between words is 500 ms, allowing strategic priming to occur. Furthermore, since the chunking rule remains the same in each block, listeners can prepare in advance about how to parse the sequences, making strategic predictions to occur more easily. Based on the knowledge about valid chunks, the semantic category of the 2^nd^ word in each chunk is fully predictable in both the same-category condition and the different-category condition. The 1^st^ word in each chunk is also predictable in the alternating-order sequences but not predictable in the random-order sequences. Since the alternating-order sequences and the random-order sequences are mixed, predictability is generally lower for the 1^st^ word than for the 2^nd^ word in each chunk. Therefore, for strategic predictions, the predictability of words correlates with the chunk structure. This kind of strategic predictions, however, is based on rule-based chunking instead of semantic relatedness stored in long-term memory.

### Rule-based Chunking Hypothesis

The rule-based chunking hypothesis posits that low-frequency neural activity reflects the grouping of words into chunks, e.g., phrases and sentences. The hypothesis is motivated by linguistic research on syntax (Chomsky, 1957) and psychological evidence for the mental representations of chunks (Miller, 1956). Psycholinguistic studies have provided evidence that the mental representation of speech is organized in the units of clauses and sentences. For example, after listening to a long sentence, words from the immediate clause can be better recalled than words from previous clauses (Jarvella, 1971; Caplan, 1972). Furthermore, if a click is presented during speech, the perceived timing of the click is attracted towards major syntactic boundaries (Fodor and Bever, 1965; Garrett et al., 1966). In terms of the neural basis, fMRI studies have demonstrated that distributed brain areas are involved in grouping words into chunks (Friederici et al., 2000; Lerner et al., 2011; Pallier et al., 2011; Bulut et al., 2017). MEG and EEG studies have suggested that low-frequency neural activity tracks linguistic structures (Ding et al., 2016; Meyer et al., 2016; Martin and Doumas, 2017; Meyer and Gumbert, 2018). Recent work suggests that animals can also parse motor sequences into hierarchically organized chunks (Geddes et al., 2018; Jiang et al., 2018).

Neural tracking of linguistic structures strongly depends on the task, demonstrating an active chunking process. Neural tracking of multisyllabic words and multi-word chunks is largely abolished during sleep (Makov et al., 2017) or when the listeners are distracted by competing sensory stimuli (Ding et al., 2018). Here, it is further demonstrated that neural tracking of a structurally ambiguous sequence relies on the chunking rule. Similar findings have been obtained when listening to a sequence of pure tones. When listeners imagine that an isochronous tone sequence is divided into groups of 2 or groups of 3, the neural responses to the tone sequence track not only individual tones but also the imagined groups (Nozaradan et al., 2011).

The chunk-rate response is consistent with the rule-based chunking model. However, it may reflect the actual chunking process or downstream processes building on the multi-word chunks. After the chunk structure is parsed, the listener could synchronize their attention and predictions to the sequence. Previous studies have suggested that entrained neural oscillations may reflect both sequence parsing (Ding et al., 2016; Kösem et al., 2016; Meyer et al., 2016; Wang et al., 2017; Meyer and Gumbert, 2018) and temporal attention/prediction (Morillon and Baillet, 2017; Jin et al., 2018; Rimmele et al., 2018), and could causally modulate speech perception (Kösem et al., 2018; Riecke et al., 2018; Zoefel et al., 2018). The current results cannot distinguish which chunk-related process drives the chunk-rate response. What can, however, be concluded here is that the chunk-rate response cannot be driven by properties of individual words, and it can only occur after the brain parses a sequence into chunks. Thus, the current study and previous studies (Ding et al., 2016) provide strong support to notion that the brain can construct superordinate linguistic representations based on either long-term syntactic rules or temporary rules learned in an experiment.

## Methods

### Participants

Thirty-two participants took part in the study (19–27 years old; mean age, 22 years; 50% female). Sixteen participants took part in Experiment 1 and the other sixteen participants took part in Experiment 2. All participants were right-handed, with no self-reported hearing loss or neurological disorders. The experimental procedures were approved by the Institutional Review Board of the Zhejiang University Interdisciplinary Center for Social Sciences and the Ethics Committee of Peking University. The participants provided written consent and were paid.

### Speech Materials

Experiment 1 presented sequences of nouns and Experiment 2 presented both noun sequences and sentence sequences. All words were disyllabic words in mandarin Chinese and each syllable was a morpheme. For the noun sequences, each word was selected from a pool of 240 disyllabic concrete nouns. These concrete nouns equally divided into 2 categories, i.e., living (L) and nonliving (N) things. Living things further divided into 2 subcategories, i.e., animals (*N* = 60; e.g., monkey, panda) and plants (*N* = 60; e.g., tulip, strawberry). Nonliving things also further divided into 2 subcategories, i.e., small manipulatable objects (*N* = 60; e.g., teacup, toothbrush) and large non-manipulatable objects (*N* = 60; e.g., playground, hotel). In each noun sequence, all living nouns were randomly drawn from a subcategory, i.e., all being animals or plants, and all nonliving nouns were also randomly drawn from a subcategory, i.e., all being manipulatable or non-manipulatable objects. Details about how the nouns constructed noun sequences were provided in the *Sequence Structure* section.

In some conditions in Experiment 2, additional 30 disyllabic abstract nouns were used to create outliers (e.g., honor, spirit). For the sentence condition in Experiment 2, 80 sentences were constructed. Each sentence had 4 syllables, with the first 2 syllables constructing a noun (or a common noun phrase) and last 2 syllables constructing a verb or (or a common verb phrase). In the following, to simplify the discussion, we refer to all the 2-syllable units as words.

For both the noun sequences and sentences sequences, each disyllabic word was independently synthesized by the iFLYTEK synthesizer (http://peiyin.xunfei.cn/; female voice, Xiaoying). All disyllabic words were adjusted to the same intensity and the same duration, i.e., 500 ms, following the procedure in Ding et al. 2016. Within a word, no additional control was applied to the intensity and duration of individual syllables and coarticulation could exist between these syllables. Compared with speech materials in which each syllable was independently synthesized, the disyllabic words synthesized as a whole sounded more natural.

When constructing sequences, the synthesized disyllabic words were directly concatenated, without any additional pause in between. Therefore, words are isochronously presented at 2 Hz. For speech stimuli generated according to this procedure, each disyllabic word was an acoustically independent unit and larger chunks consisting of multiple words had no acoustically defined boundaries.

### Sequence Structure

In Experiment 1, pairs of nouns constructed chunks and chunks further constructed sequences. The experiment compared 2 conditions in which the chunks were constructed based on different rules. For the same-category condition, the 2 nouns in each chunk belonged to the same category. For the different-category condition, however, the 2 nouns in each chunk were from different categories. Since the study only considered 2 categories of words, there were 2 valid chunks in the same-category condition, i.e., LL and NN, and 2 valid chunks in the different-category, i.e., NL and LN. Each chunk is 1 s in duration.

Each sequence consisted of 12 chunks and therefore was 12 s in duration. In each sequence, the 2 valid chunks were concatenated in either an alternating order or a random order (Fig. 2A). The alternating-order sequence in each condition had a fixed structure, repeating a 4-words unit 6 times, i.e., NNLL for the same-category condition and NLLN for the different-category condition. In each random-order sequence, every chunk was randomly and independently chosen from the 2 valid chunks. After the category of each word was determined and the actual words were filled in. Each word was randomly drawn from a pool of 60 words (see *Speech Materials*), with the additional constraint that no word repeated in a sequence.

Experiment 2 considered 3 kinds of sequences, all of which consisted of 24 disyllabic words and was 12 s in duration. One kind of sequence was the same as the alternating-order sequence constructed by same-category chunks in Experiment 1. In the 2^nd^ kind of sequence, each noun was randomly chosen from the living and nonliving nouns, without any chunk structure. The 3^rd^ kind of sequence was constructed by the 4-syllable sentences.

### Experimental procedures and tasks

#### Experiment 1

The same-category condition and the different-category were presented in separate blocks and the order of the 2 blocks was counterbalanced across participants. In each condition, 30 alternating-order sequences and 30 random-order sequences were mixed and presented in a random order. In 8 alternating-order sequences and 8 random-order sequences, the living noun in one chunk was switched with the nonliving noun in another chunk so that the 2 chunks were no longer valid (Fig. 2B). These 16 sequences with invalid chunks were outlier sequences. The outlier sequences (*N* = 16) and normal sequences (*N* = 44) were mixed and presented in a random order. However, only the normal sequences are involved in the neural response analysis. The participants had a rest after listening to 30 sequences.

Before each condition, instructions were given about the chunk structures. During the experiment, participants were asked to mentally segment the sequences into 2-word chunks and judge whether all the chunks were valid chunks. In other words, they had to distinguish normal and outlier sequences and indicate their decisions by pressing different keys at the end of each sequence. After the key press, the next sequence was presented after a silent interval randomized between 1 and 2 s (uniform distribution).

At the beginning of the experiment, participants were familiarized with all the synthesized words. In the familiarization session, after hearing a word, the participants pressed a key to see the word on a screen. Then, the participants could press one key to hear the word again or press another key to hear the next word. After this familiarization session, the participants learned the sequence structure and listened to 2 normal sequences and 2 outlier sequences, which were presented in random order. They had to tell the experimenter whether they heard outliers and what the outlier chunks were. Next, a practice session was given which was the same as the MEG experiment, except that it was ended after the participants made 4 correct responses in 5 consecutive sequences. In the MEG experiment, the participants made correct responses in 85 ± 2% and 86 ± 2% of sequences for the same-category condition and different-category condition, respectively (mean ± SEM across participants).

#### Experiment 2

The experiment consisted of 5 conditions that were presented in separate blocks. In 3 conditions, participants performed different tasks while listening to the alternating-order sequence of same-category chunks. One task was a chunk-level task that had to detect weather an invalid chunk appeared at a random position in the sequence. The 2^nd^ task was a word-level task, detecting if a concrete noun at a random position was replaced by an abstract noun. The 3^rd^ task was an auditory task, detecting if the speaker’s voice was changed for a word at a random position of the sequence. The voice change was implemented by the change-gender function in Praat (Boersma, 2006). The participants made correct responses in 80 ± 5%, 91 ± 2% and 96 ± 2% of sequences during the chunk-level task, word-level task and auditory task, respectively.

The other 2 conditions presented random word sequences and sentence sequences, respectively. The participants performed a word-level task in these 2 conditions, detecting whether a word was replaced by an abstract noun. The participants made correct responses in 95 ± 1% and 88 ± 2% of sequences for the random word sequences and sentence sequences, respectively.

After listening to a sequence, participants pressed different keys to indicate whether they detected an outlier or not. After the key press, the next sequence was presented after a silent interval randomized between 1 and 2 s (uniform distribution). Each condition consisted of 20 normal sequences and 5 outlier sequences, which were mixed and presented in a random order. Only the normal sequences were involved in the neural data analysis. All 5 conditions were presented in a random order, with the constraint that the 3 conditions using alternating-order chunk sequences are next to each other and the random word and sentence conditions are also next to each other. Participants were informed of the task before each stimulus condition.

Before the MEG recording section, participants were familiarized with all the synthesized words using the same procedure in Experiment 1. Participants were also familiarized with the task before each condition by listening to 1 normal sequence and 1 outlier sequence.

### Data Acquisition

Neuromagnetic responses were recorded using a 306-sensor whole-head MEG system (Elekta-Neuromag, Helsinki, Finland) at Peking University, sampled at 1 kHz. The system had 102 magnetometer and 204 planar gradiometers. Four MEG-compatible electrodes were used to record EOG at 1000 Hz. Two electrodes were placed at the left and right temples and their difference was the horizontal EOG (right minus left). Another 2 electrodes were placed above and below the right eye and their difference was the vertical EOG (upper minus lower). To remove ocular artifacts in MEG, the horizontal and vertical EOG were regressed out from the MEG recordings using the least-squares method.

Four head position indicator (HPI) coils were used to measure the head position inside MEG. The positions of 3 anatomical landmarks (nasion, left, and right pre-auricular points), the 4 HPI coils, and at least 200 points on the scalp were also digitized before experiment. For MEG source localization purposes, structural Magnetic Resonance Imaging (MRI) data were collected from all participants using a Siemens Magnetom Prisma 3-T MRI system (Siemens Medical Solutions, Erlangen, Germany) at Peking University. A 3-D magnetization-prepared rapid gradient echo T1-weighted sequence was used to obtain 1 × 1 × 1 mm^3^ resolution anatomical images.

### Data Processing

Temporal Signal Space Separation (tSSS) was used to remove the external interference from MEG signals (Taulu and Hari, 2009). Since the current study only focused on responses at 1 and 2 Hz, the MEG signals were bandpass filtered between 0.5 and 3 Hz using a linear-phase finite impulse response (FIR) filter, and downsampled at 20 Hz. The response during each sequence was extracted and was referred to as a trial. The MEG signals were further denoised using a semi-blind source separation technique, the Denoising Source Separation (DSS). The DSS was a linear transform that decomposed multi-sensor MEG signals into components (de Cheveigné and Simon, 2008). The bias function of the DSS was chosen as the response averaged over trials within each condition. A common DSS for all conditions was derived based on the response covariance matrices averaged over conditions. The first 6 DSS components were retained and transformed back to the sensor space for further analysis. This DSS procedure was commonly used to extract cortical responses entrained to speech (Ding et al., 2016; Zhang and Ding, 2017).

### Source Localization

The MEG responses averaged over trials were mapped into source space using cortex constrained minimum norm estimate (MNE) (Hämäläinen and Ilmoniemi, 1994), implemented in the Brainstorm software (Tadel et al., 2011). The T1-weighted MRI images were used to extract the brain volume, cortex surface, and innermost skull surface using the Freesurfer software (http://surfer.nmr.mgh.harvard.edu/). In the MRI images, the 3 anatomical landmarks (nasion, left, and right pre-auricular points) were marked manually. Both three anatomical landmarks and digitized head points were used to align the MRI images with MEG sensor array. The forward MEG model was derived based on the overlapping sphere model (Huang et al., 1999). The identity matrix was used as noise covariance. Source-space activation was measured by the dynamic statistical parametric map (dSPM) (Dale et al., 2000) and the value was in arbitrary unit (a.u.). Individual source-space responses, consisting of 15,002 elementary dipoles over the cortex, was rescaled to the ICBM 152 brain template (Fonov et al., 2011) for further analyses.

### Frequency-domain analysis

In the frequency-domain analysis, to avoid the onset response, the response during the first second of each trial were removed. Consequently, the neural response was 11 s in duration for each trial. The average of all trials was transformed into the frequency domain using the Discrete Fourier Transform (DFT) without any additional smoothing window. The frequency resolution of the DFT analysis was 1/11 Hz. If the complex-valued DFT coefficient at frequency *f* was denoted as *X*(*f*), the response power and phase were |*X*(*f*)|^2^ and ∠*X*(*f*) respectively. For the response power analysis, responses from the 2 collocated gradiometers were always averaged. Additionally, normalized power was calculated to compensate the baseline response power. The normalized power at frequency *f* was the difference between the power at *f* and the power averaged over 4 neighboring frequency bins (2 bins on each side). For the phase analysis, all magnetometers and gradiometers were separately analyzed. The circular mean was used to average the neural response phase over participants or sensors. The circular phase coherence was used to measure the spread of response phase across participants (Fisher, 1993).

### Statistical tests

All tests were based on bias-corrected and accelerated bootstrap (Efron and Tibshirani, 1994). In the bootstrap procedure, all participants were resampled with replacement 10000 times. All comparisons in this study were paired comparisons. For one-sided comparison of response power, if the response power in one condition was greater than the other condition in *A*% of the resampled data, the significance level is (100*A* + 1)/10001. For two-sided comparisons, if the response power was greater in one condition for *A*% of the resampled data, the significance level is (200*A* + 1)/10001.

#### Spectral peak

The statistical significance of a spectral peak at frequency *f* was tested by comparing the response power at *f* with the power averaged over 4 neighboring frequency bins (2 bins on each side, one-sided comparison). The significance test was only applied to the response power at the chunk and word rates. A false discovery rate (FDR) correction was applied to these two frequencies.

#### Power difference between conditions or hemispheres

A two-sided test was used to compare the normalized power between conditions. To characterize response lateralization in the sensor space, normalized response power was averaged over the left and right hemispheres respectively (96 gradiometers in each hemisphere).

#### Response phase

A two-sided test was used to compare the response phase difference between conditions. The *A*% confidence interval of the phase difference was measured by the smallest angle that could cover *A*% of the 10000 resampled phase difference. If the confidence interval did not include 0° or 180°, the response phase significantly deviated from 0° or 180° (significance level 1 − *A*%).

### Model Simulations

The lexical property model and the semantic relatedness model were based on lexical features. For the simple model illustrated in Fig. 2, only 2 features were considered, i.e., is-living or is-nonliving. Each feature took a binary value, i.e., 1 for yes and 0 for no. For the realistic model illustrated in Supplementary Fig. 2, 300 feature dimensions were used and each feature was coded by a real number. The features were derived based on the word2vec model (Bengio et al., 2003) and the model was trained based on large copora (the ‘combination’ corpora) (Li et al., 2018). The word2vec model is built on recurrent neural networks that do not consider phrasal structures.

In model simulations, each feature dimension was denoted by a pulse sequence. A pulse was placed at the onset of each word and its amplitude was the feature value (Supplementary Fig. 1). For the word2vec model, the amplitude could be negative. For the lexical property model, the neural response was simulated by convolving the feature pulse sequence with a response function, which was a 500-ms duration Gaussian window. For the lexical property model, the neural response to each feature was independently simulated and transformed into frequency domain. For the simple model in Fig. 2, the 2 feature dimensions were shown separately. For the word2vec model that used 300 feature dimensions, the power spectrum was averaged over feature dimensions (Frank and Yang, 2018).

For the semantic similarity model, the similarity between feature vectors was measured by the Euclidean distance (Supplementary Fig. 1). A pulse sequence denoting the Euclidean distance between the current word and the previous word was used to simulate the neural response. Based on this method, if neighboring words were represented by similar feature vectors, their distance would be small and consequently the neural response amplitude would be small, consistent with the neural adaptation effect.

The rule-based chunking model predicted a consistent change of neural activity within the duration of a chunk, but had no specific assumptions about the waveform of the neural response. Here, to facilitate the comparison with the semantic relatedness model, it is further assumed that the response was stronger at the chunk onset. The rule-based chunking model was also simulated by convolving the response function with a pulse sequence. In the pulse sequence, a pulse was placed at the word onset. The pulse amplitude was 1 for the first word in the chunk and was 0.5 for the second word in the chunk.

## Acknowledgement

We thank Cheng Cheng for excellent assistant in data collection, David Poeppel, Lucia Melloni, Lang Qin, Jiajie Zou, and Cheng Luo for thoughtful comments on previous versions of the manuscript. Work supported by National Natural Science Foundation of China 31771248 (ND) and Zhejiang Provincial Natural Science Foundation of China LR16C090002 (ND).

---

**Supplementary Figure 1.**
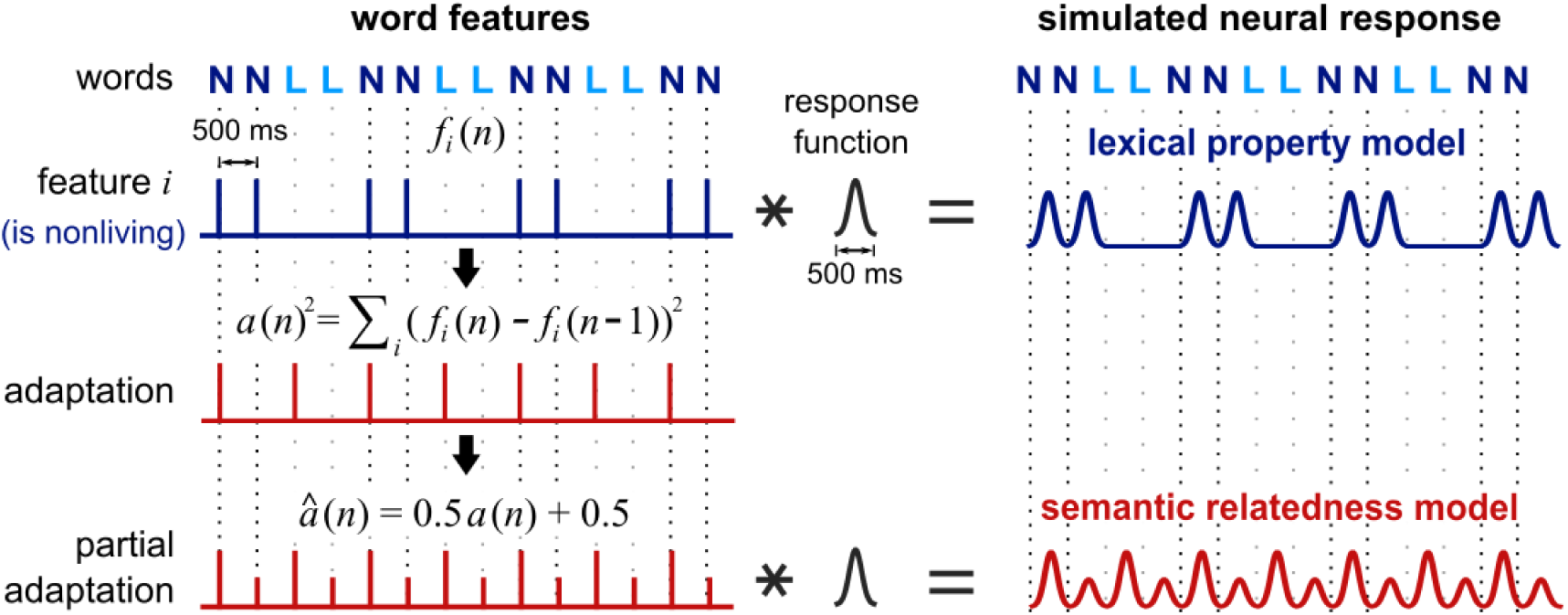
Procedures to simulate the lexical property model and the semantic relatedness model. The left panel illustrates the representation of lexical features. Each feature dimension is represented by a pulse sequence, with a pulse being placed at the onset of each word and the amplitude of the pulse being modulated by the word feature. One feature dimension is illustrated here and it is a binary feature denoting nonliving things. For the models using word2vec features, each feature is coded by a real number instead of the binary numbers shown in this illustration. Neural response predicted by the lexical property model is simply the feature sequence convolving a response function, which is a 500-ms Gaussian window. For the semantic relatedness model, the Euclidean distance is used to measure semantic similarity between feature vectors. Since the neural response does not completely disappear even the same word repeats, we provide a DC offset to the Euclidean distance sequence, i.e., the partial adaptation model in the figure.

**Supplementary Figure 2.**
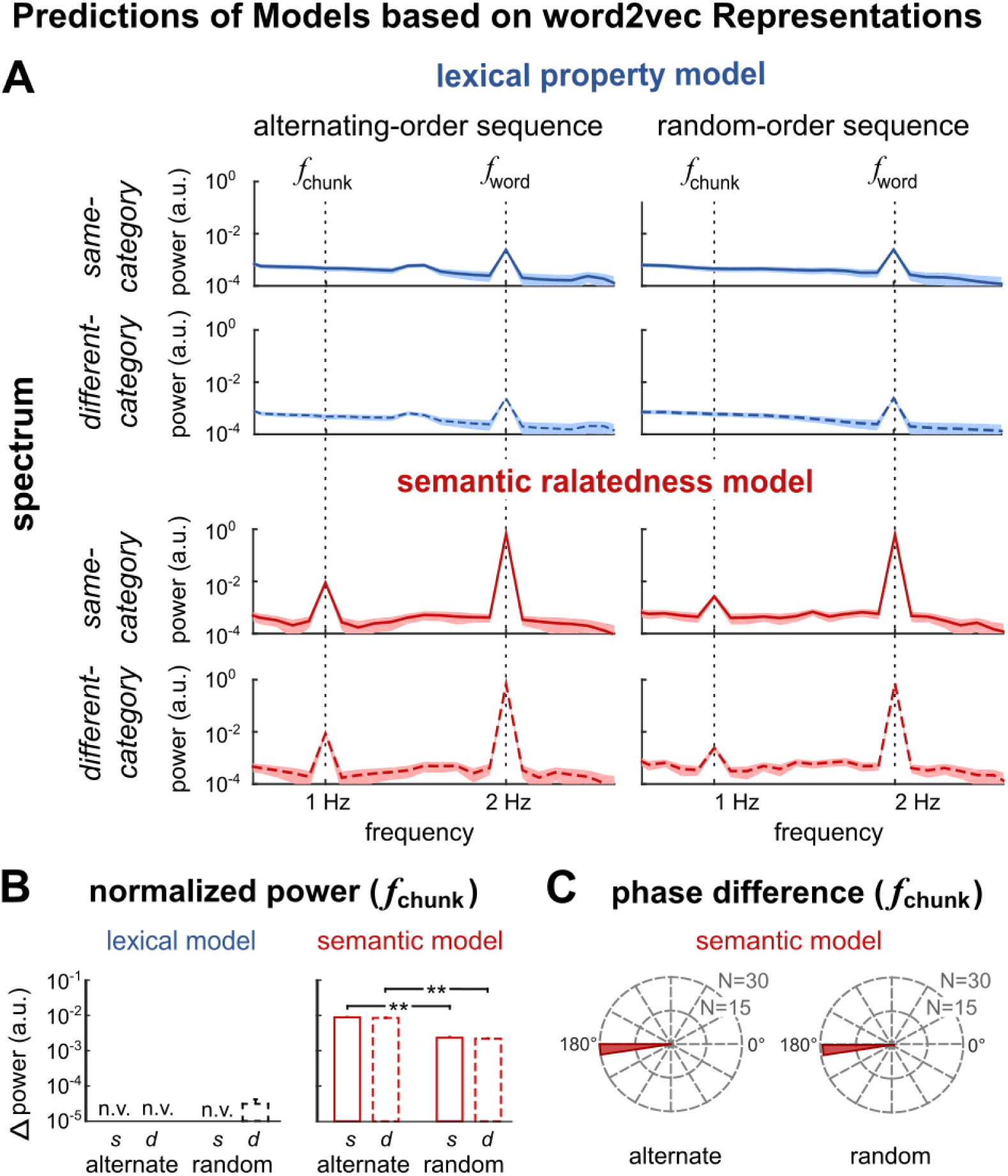
Model simulations based on word representations learned from natural language. The 300-dimensional vectorial representation of each word is learned from large corpus using the word2vec model (Bengio, et al., 2003; Li et al., 2018). Unlike the simulations in Fig. 2, the 300-dimentional features here are purely data driven, not necessarily corresponding to specific semantic categories, e.g., living and nonliving things. For lexical property model, the neural responses to all 300 features were averaged in frequency domain. For semantic relatedness model, the semantic relatedness between neighboring words is measured by the Euclidean distance between the 300-dimentional representations. The rule-based model does not depend on word representations and therefore is not plotted here. (A) Response spectrum. The semantic relatedness model, but not the lexical property model, generates a chunk-rate response, consistent with the results in Fig. 2. (B) The normalized chunk-rate response power. For the lexical property model, the normalized power has negative values (n.v.) or is close to zero. The semantic relatedness model predicts a difference in response power between alternating- and random-order sequences, consistent with the results in Fig. 2. (C) The semantic relatedness model predicts a 180° phase difference between same- and different-category conditions, consistent with the results in Fig. 2. The phase difference is not shown for the lexical property model since it predicts no significant chunk-rate response. * P <0.05, ** P < 0.01

